# Antagonistic role of the BTB-zinc finger transcription factors Chinmo and Broad-Complex in the juvenile/pupal transition and in growth control

**DOI:** 10.1101/2022.11.17.516883

**Authors:** Sílvia Chafino, Panagiotis Giannios, Jordi Casanova, David Martín, Xavier Franch-Marro

## Abstract

During development, the growing organism transits through a series of temporally regulated morphological stages to generate the adult form. In humans, for example, development progresses from childhood through to puberty and then to adulthood, when sexual maturity is attained. Similarly, in holometabolous insects, immature juveniles transit to the adult form through an intermediate pupal stage when larval tissues are eliminated and the imaginal progenitor cells form the adult structures. The identity of the larval, pupal and adult stages depends on the sequential expression of the transcription factors *chinmo, Br-C* and *E93*. However, how these transcription factors determine temporal identity in developing tissues is poorly understood. Here we report on the role of the larval specifier *chinmo* in larval and adult progenitor cells during fly development. Interestingly, *chinmo* promotes growth in larval and imaginal tissues in a Br-C-independent and -dependent manner, respectively. In addition, we found that the absence of *chinmo* during metamorphosis is critical for proper adult differentiation. Importantly, we also provide evidence that, in contrast to the well-known role of *chinmo* as a pro-oncogene, Br-C and E93 act as tumour suppressors. Finally, we reveal that the function of *chinmo* as a juvenile specifier is conserved in hemimetabolous insects as its homolog has a similar role in *Blatella germanica*. Taken together, our results suggest that the sequential expression of the transcription factors Chinmo, Br-C and E93 during larva, pupa an adult respectively, coordinate the formation of the different organs that constitute the adut organism.

## Introduction

Animal development passes through various stages characterised by distinct morphological and molecular changes. In humans, for instance, development continues from birth through to childhood and puberty to give rise to the adult form. As in many animals, in holometabolous insects such as *Drosophila melanogaster*, the developmental stages are sharply defined: embryogenesis gives rise to the larva, a juvenile stage, which, upon different rounds of growth and moulting, brings about a new stage structure, the pupa, when most of the larval cells die and the adult progenitor cells (imaginal cells) develop to generate the adult organism. The regulation of stage-specific differences is mediated by the action of two major developmental hormones, the steroid 20-hydroxyecdysone and the terpenoid juvenile hormone (Hiruma and Kaneko, 2013; Jindra et al., 2013; Truman, 2019; Truman and Riddiford, 2007, 2002; Yamanaka et al., 2013). Both hormones exert this precise developmental control by regulating the expression of three critical genes that encode for the stage-identity factors that compose the Metamorphic Gene Network (MGN): the C2H2 zinc finger type factor *Krüppel-homolog 1* (*Kr-h1*), the the helix turn–helix *Ecdysone inducible protein 93F* (*E93*), and *Broad-complex* (*Br-C*; also known as broad), a member of the bric-a-brac-tramtrack-broad family (Martín et al., 2021).

The deployment of the pupal-specific genetic program is controlled by the expression of *Br-C* at the larval–pupal transition (Truman, 2019; Zhou and Riddiford, 2002). Upon the formation of the pupa, hormone signalling triggers the expression of the helix-turn-helix factor *E93*, whose product represses *Br-C* expression and directs the formation of the final differentiated adult structures (Chafino et al., 2019; Martín et al., 2021; Ureña et al., 2014). While it is firmly established that *Br-C* and *E93* are the stage-specifying genes for the pupal and adult states, the nature of the larval specifying gene has been elusive. To date, larval identity has been attributed to Kr-h1, which is present during the larval period and represses *Br-C* and *E93* expression during this period (Huang et al., 2011; Ureña et al., 2016). However, although Kr-h1 is undoubtedly critical for maintaining the larval state, evidence has shown that this factor cannot be considered the larval specifier *per se*. For example, depletion of *Kr-h1* in *Drosophila* does not prevent normal larval development nor a timely transition to the pupa (Beck et al., 2004; Pecasse et al., 2000). In this regard, the product of *chronologically inappropriate morphogenesis* (*chinmo*) gene, another member of the BTB family of transcription factors, has been recently proposed to be responsible for larval identity in *Drosophila* (Truman and Riddiford, 2022).

First isolated based on its requirement for the temporal identity of mushroom body neurons (Zhu et al., 2006), the identification of Chinmo as a more general larval specifier has provided invaluable insights into the molecular mechanisms underlying the control of juvenile identity. Yet, little is known about how this factor exerts its function along with Br-C and E93. Moreover, given that holometabolous insects comprise both larval tissues and pools of adult progenitor cells, a central issue in the understanding of how larval identity is controlled is how larval and adult progenitor cells respond differentially to the same set of temporal transcription factors. Furthermore, in the sequential activation of *chinmo, Br-C* and *E93*, the extent of the activity directly attributable to each transcription factor or to their mutual repression is still unclear.

Here we confirm the role of *chinmo* as larval specifier in larval and adult progenitor cells and establish its regulatory interactions with the other temporal specifiers. We also examine how the temporal sequence of Chinmo and Br-C differently affects with the genetic program that establishes larval vs. imaginal identity. Thus, we found that Chinmo controls larval development of larval and imaginal tissues in a Br-C-independent and -dependent manner, respectively. According to these data, and in the context of the MGN, we also show that *chinmo* absence is critical for the transition from larva to pupa and then to adult, as it acts as a repressor of both *Br-C* and *E93*. In addition, we report that the *chinmo* homologue has a similar role in the cockroach *Blatella gemanica*, thereby indicating that its function as a juvenile specifier precedes the hemimetabolous/holometabolous split. Finally, we show that in contrast to the well characterized role of *chinmo* as pro-oncogene, the *Br-C* pupal and E93 adult specifiers have those of tumour suppressor genes. These characteristics are maintained besides insects and may account for the different role of some human BTB zinc-finger transcription factors in tumorigenesis.

## Results and Discussion

### *chinmo* is expressed throughout larval stages and is required in larval and imaginal tissues

Examination of *chinmo* expression revealed that it is expressed during embryogenesis and early larval development and that it is strongly downregulated from L3 (Figure 1A). Immunostaining analysis in imaginal and larval tissues confirmed the presence of Chinmo in L1 and L2 stages and its disappearance in late L3 (Figure 1B and C), an expression profile that is in agreement with previous studies (Narbonne-Reveau and Maurange, 2019; Truman and Riddiford, 2022). We next addressed its functional requirement by knocking down this factor with an RNAi transgene controlled by the ubiquitous *ActGal4* driver. *chinmo*-depleted animals showed developmental arrest at the end of the first instar larval stage presenting a tanned cuticle clearly reminiscent to that of the pupa (Figure 1D). Consistent with the phenotype, we found that arrested *chinmo*-depleted larvae precociously expressed pupal cuticle genes while blocked larval specific genes activation (Figure 1E). These results confirm that *chinmo* is required for normal progression of the organism during the larval period, as proposed by Truman and Riddiford (Truman and Riddiford, 2022).

**Figure 1.**
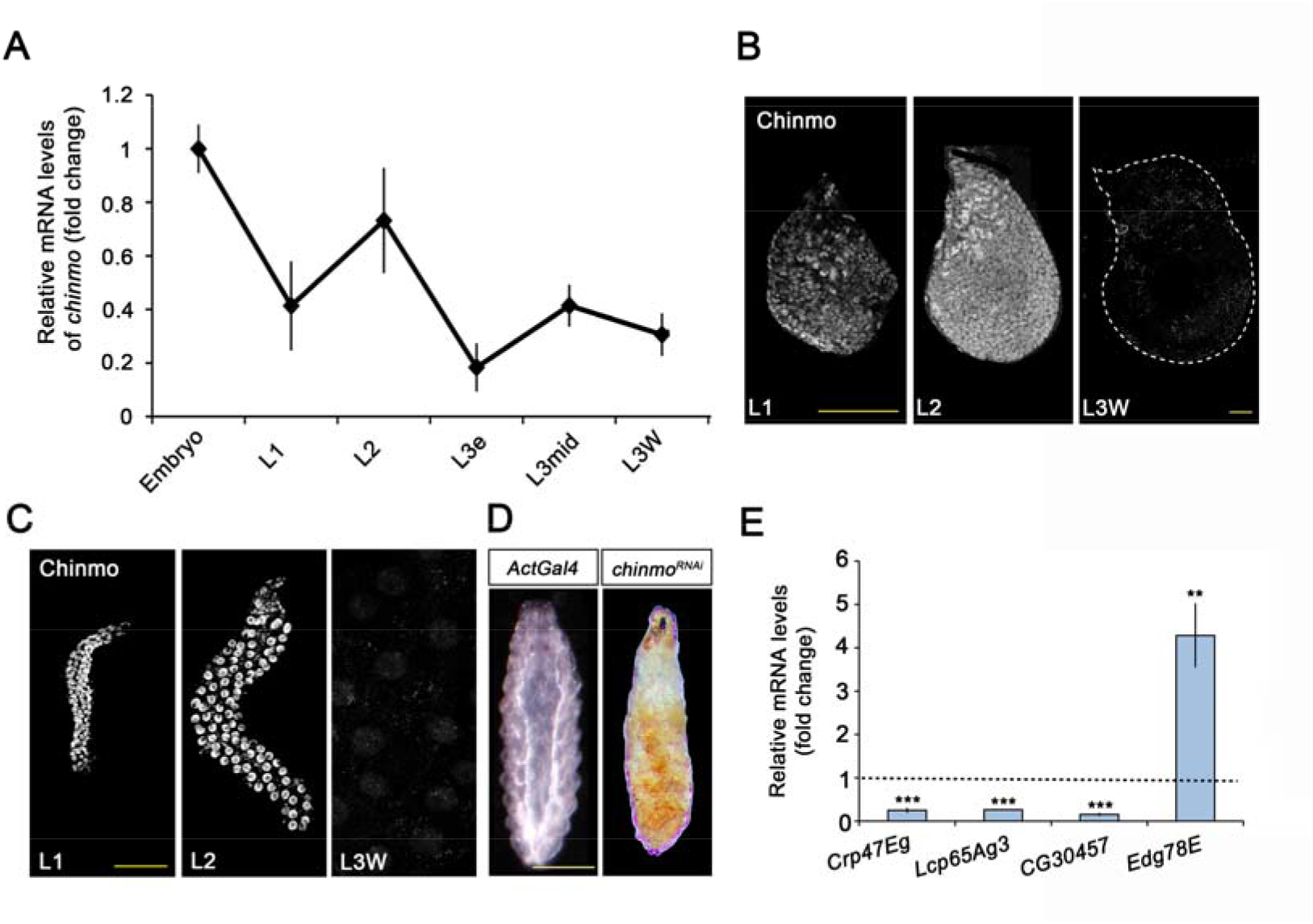
Chinmo is expressed during early larval stages and is essential for proper larval development. **(A)** *chinmo* mRNA levels measured by qRT-PCR from embryo to the wandering stage of L3 (L3W). Transcript abundance values were normalised against the *Rpl32* transcript. Fold changes were relative to the expression of embryo, arbitrarily set to 1. Error bars indicate the SEM (n = 3). (B-C) Chinmo protein levels in the wing disc (B) and SGs (C) of larval L1, L2 and L3W stages. (D) Compared with the control (*ActGal4*), overexpression of *UAS chinmo*^*RNAi*^ in the whole body induced developmental arrest at the L1 stage. Scale bars represent 50 µm (B and C) and 0.5 mm (D). (E) Relative expression of larval-specific (*Crp47Eg, Lcp65Ag3 and CG30457*) and pupal-specific genes (Edg78E) in *UASchinmo*^*RNAi*^ L1 larvae measured by qRT-PCR. Transcript abundance values were normalised against the *Rpl32* transcript. Fold changes were relative to the expression in control larvae, arbitrarily set to 1 (dashed black line). Error bars indicate the SEM (n = 3). Statistical significance was calculated using t test ( ∗ ∗ ∗*p ≤ 0*.*001*; *\*\*p ≤ 0*.*005*). **Figure 1—Source Data 1** **Numerical data for Figure 1A and E**.

Since *Drosophila* larva consists of a combination of larval and imaginal tissues, we then analysed the contribution of *chinmo* to the development of these two types of tissues. Regarding the former, *chinmo* was selectively depleted in the salivary glands (SGs) using the *forkhead* (*fkh*) driver (*fkhGal4*), which is active in this tissue from embryogenesis onwards. The SG is a secretory organ that develops from embryonic epithelial placodes (Abrams et al., 2003; Bradley et al., 2001; Camelo and Luschnig, 2021) and increases dramatically in size by cell endoreplication during the larval period (Edgar et al., 2014; Zielke et al., 2013). This tissue is responsible for producing glycosylated mucin for the lubrication of food during the larval period (Costantino et al., 2008; Farkaš et al., 2014; Riddiford, 1993; Syed et al., 2008) and for synthesising glue proteins for the attachment of the pupa to a solid surface at the onset of metamorphosis (Andres et al., 1993; Costantino et al., 2008; Kaieda et al., 2017). As it is shown in Figure 2A, although depletion of *chinmo* in the SG did not affect the formation of this organ, it caused a dramatic decrease in normal larval development, as revealed by the strong reduction in size and DNA content of the gland cells (Figure 2B-D). Consistently, the expression levels of *salivary gland secretion* (*sgs*) genes, which encode for several components of the glue, were markedly repressed in *chinmo* depleted SG compared to control (Figure 2E).

**Figure 2.**
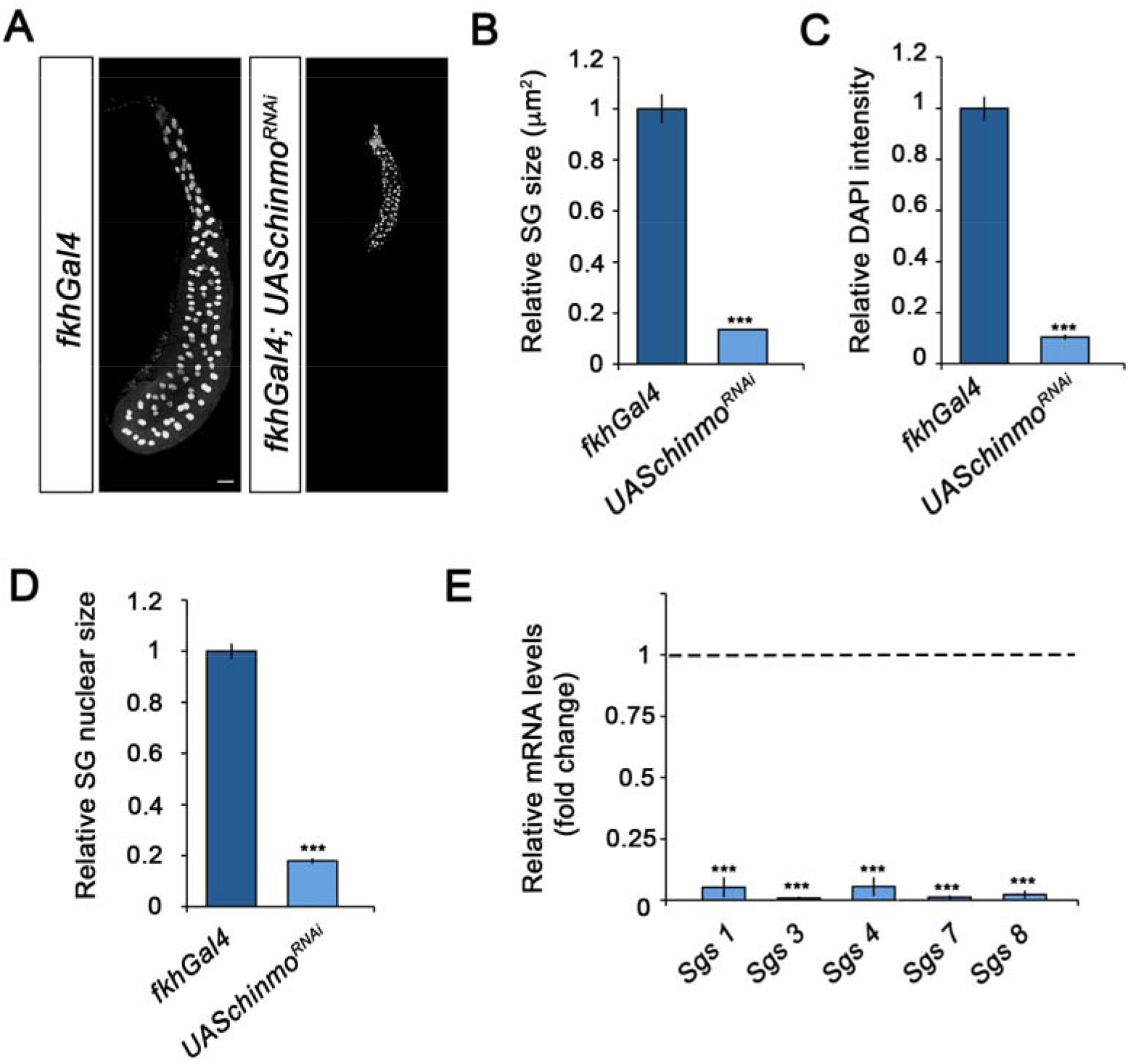
Chinmo is required for proper growth and function of the SG during larval development. (A) DAPI-staining of SGs from control (*fkhGal4*) and *UASchinmo*^*RNAi*^ larvae at L3W. Scale bar represents 50 µm. (B-D) Comparison of the relative size of SGs (n = 10 for each genotype) (B), DAPI intensity (n = 50 for each genotype) (C), and nucleic size of SGs (n = 50 for each genotype) (D) between *UASchinmo*^*RNAi*^ and control larvae at L3W. Error bars indicate the SEM (n = 5-8). (E) Relative expression of SG secretion genes in *UASchinmo*^*RNAi*^ animals measured by qRT-PCR. Transcript abundance values were normalised against the *Rpl32* transcript. Fold changes were relative to the expression in control larvae, arbitrarily set to 1 (dashed black line). Error bars indicate the SEM (n = 5-8). Statistical significance was calculated using t test (∗ ∗ ∗*p ≤ 0*.*001*). **Figure 2—Source Data 2** **Numerical data for Figure 2B-E**.

Regarding the role of *chinmo* in imaginal tissues, we knocked down this factor in wing imaginal discs from the embryonic period onwards using the *escargot* (*esg*) driver (*esgGal4*). As before, depletion of *chinmo* in the *esg* domain did not alter the specification of the disc, but strongly impeded its larval development. Thus, in late L3 wing discs only the notum, which does not express the *esgGal4* driver, was observed while the wing pouch, revealed by positive GFP signal, was strongly reduced and did not show the expression of patterning genes such as *wingless* (*wg*) and *cut* (*ct*) (Figure 3A). In line with these results, although most of the *chinmo* depleted animals arrested development as pharate adults, escapers that were able to eclose (15%) had no wings (SFigure 1). Taken together, these data show that Chinmo is required during the larval period to control the development and function of larval and imaginal tissues.

**Figure 3.**
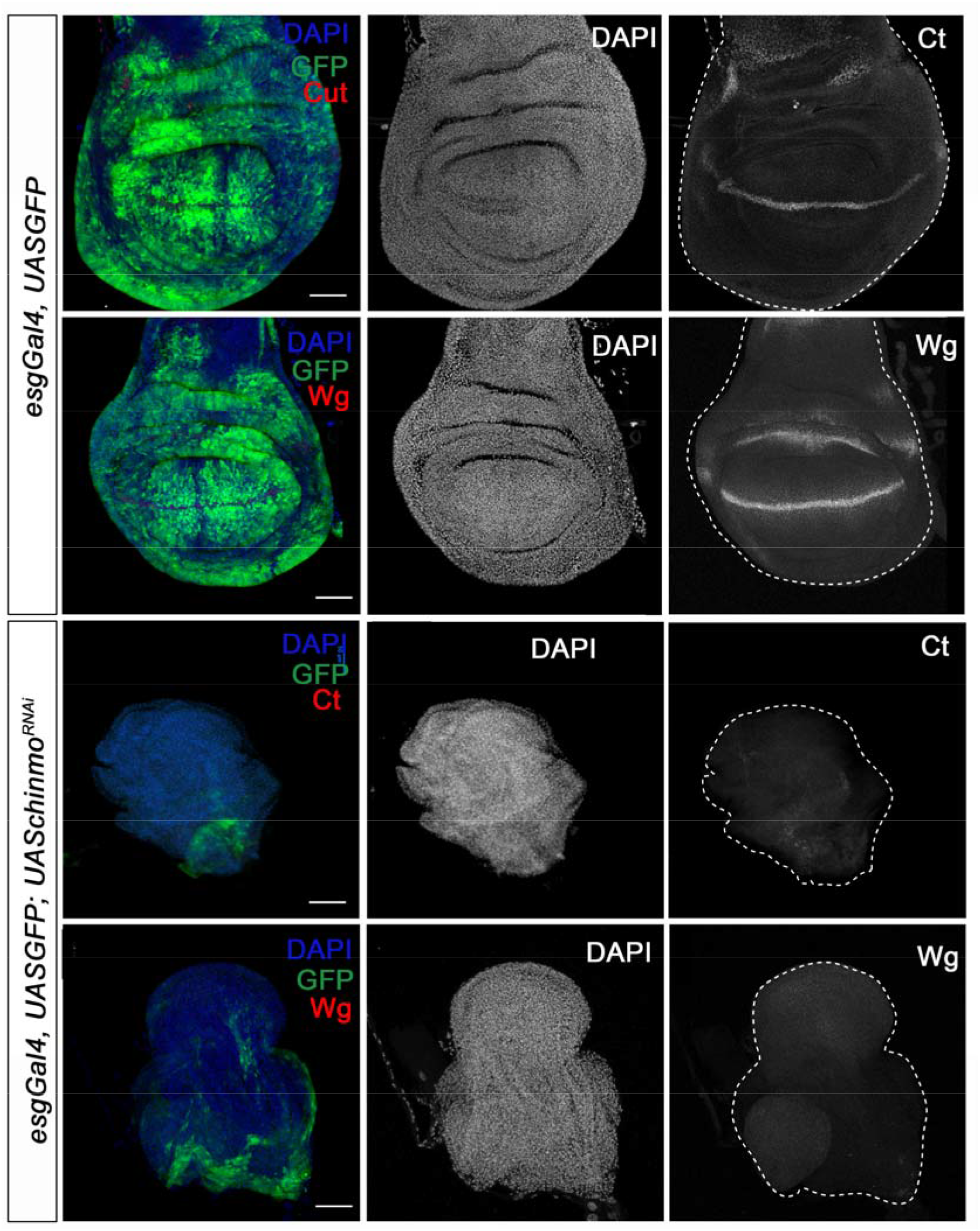
Chinmo is necessary for wing development during the larval period. Expression of Ct and Wg in wing discs of control (*esgGal4*) and *UASchinmo*^*RNAi*^ L3W larvae. Wing discs were labelled to visualise the *esg* domain (GFP in green) and nuclei (DAPI). Ct and Wg were not detected in *UASchinmo*^*RNAi*^.

### Distinct roles of Chinmo in larval and progenitor cells

A critical feature of the MGN factors is that their sequential expression is achieved through a series of regulatory interactions between them. Therefore, we next sought to characterise the regulatory interactions of Chinmo with the pupal specifier Br-C and the adult specifier E93. To this end, we measured the expression of *Br-C* and *E93* in *chinmo*-depleted SGs and wing discs. Contrary to recently published data (Truman and Riddiford, 2022), both tissues showed a significant and premature increase of Br-C protein levels as early as in L1 larvae, while no increase in E93 protein levels was detected in any tissue (Figure 4).

**Figure 4.**
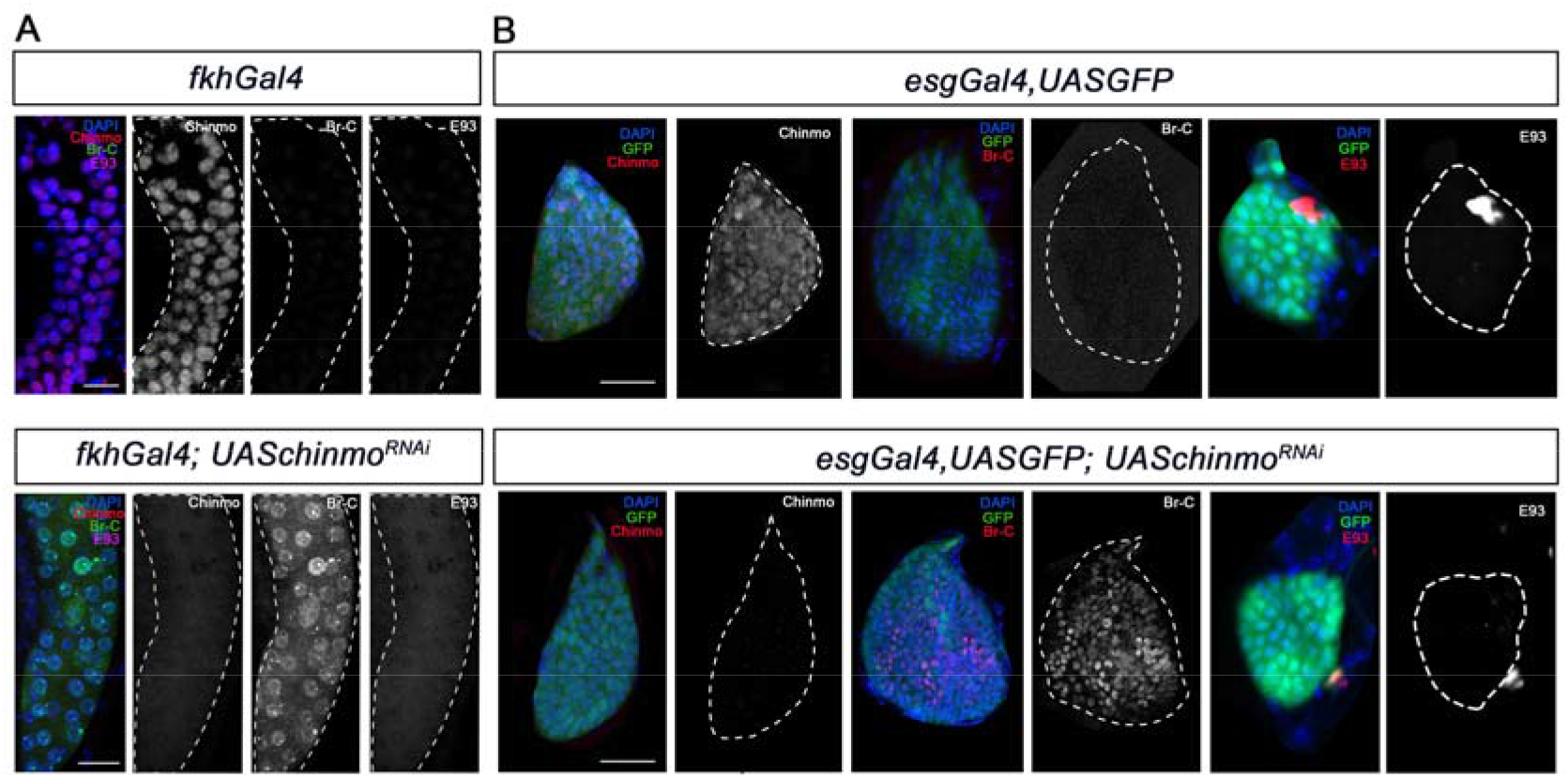
Chinmo represses *Br-C* in SGs and wing discs during early larval development. (A) Expression of chinmo, Br-C and E93 in SGs of L1 control (*fkhGal4*) and *UASchinmo*^*RNAi*^. (B) Expression of Chinmo, Br-C and E93 in wing discs of early L2 control (*esgGal4*) and *UASchinmo*^*RNAi*^. The *esg* domain is marked with GFP and all cell nucleus with DAPI. In the absence of *chinmo* only *Br-C* shows early up-regulation in both tissues. Scale bars represent 25 µm.

In view of these results, we speculated whether the impairment of larval development observed in *chinmo*-depleted animals could be the result of precocious presence of the wrong stage-identity factor, in this case, Br-C. To address this notion, we precociously expressed Br-CZ1, the main Br-C isoform expressed during imaginal larval development (Narbonne-Reveau and Maurange, 2019), in SGs and wing discs. Interestingly, as previously described ectopic expression of Br-CZ1 blocked Chinmo activation (Narbonne-Reveau and Maurange, 2019 and data not shown). As a consequence precocious upregulation of Br-C phenocopied the loss of function of *chinmo* as blocks development in both tissues (SFigure 2). This result suggests that the main function of Chinmo is to avoid the expression of the pupal specifier Br-C during the juvenile stages. To confirm this hypothesis we simultaneously depleted *chinmo* and *Br-C* in SGs and wing discs. Remarkably, whereas SGs showed the same growth impairment as that observed upon *chinmo* depletion (Figure 5A-E), the absence of *Br-C* and *chinmo* largely rescued the abnormalities of the wing discs seen upon depletion of this transcription factor. The double knock-out wing discs developed in a regular manner to reach normal size by the end of L3 and showed proper expression of patterning genes such as *wg* (Figure 5F). Taken together, our results suggest that a major regulatory function of chinmo during early larval development in adult progenitor cells is channelled through the repression of *Br-C*, while in larval tissues it appears to exert specific growth-related functions that are independent to *Br-C* repression. Thus, Chinmo ensures the expression of juvenile genes by repressing Br-C, a well known inhibitor of larval gene expression (Zhou and Riddiford, 2002). In this regard, it is tempting to speculate that Br-C might repress the early expression of critical components of signaling pathways such as wingless and EGFR, involved in wing fate specification in early larval development (Ng et al., 1996; Wang et al., 2000; Zecca and Struhl, 2002). In contrast, Chinmo seems to exert an active role promoting growth in larval tissues. This different response could be explained by the nature of the two types of tissues. While larval tissues are mainly devoted to growth during the larval period and then fated to die during the metamorphic transition, the developmental identity of the imaginal cells is modified along the larva–pupa–adult temporal axis to give rise to the adult structures. In fact, it has already been shown that the other members of the MGN exert different functions depending on the type of tissue. For example, while Br-C is necessary for the degeneration of the SG during the onset of the pupal period (Jiang et al., 2000), it is critical for the correct eversion of the wing disc and for the temporary G2 arrest that synchronizes the cell cycle in the wing epithelium during early pupa wing elongation (Guo et al., 2016). Likewise, E93 is necessary to activate autophagy for elimination of mushroom body neuroblasts in late pupae (Pahl et al., 2019), whereas it controls the terminal adult differentiation of the wing during the same period (Ureña et al., 2016; Uyehara et al., 2017).

**Figure 5.**
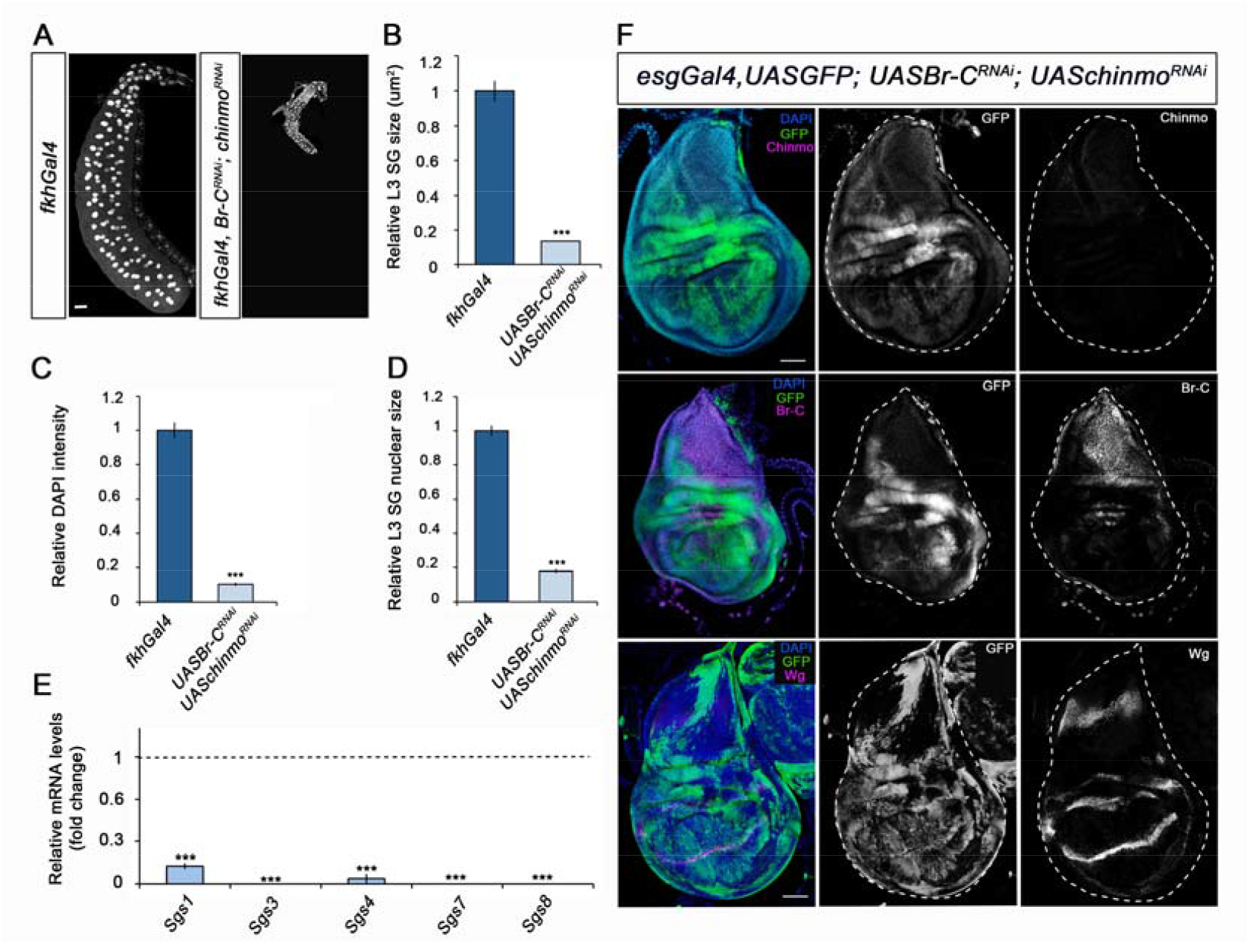
Different requirement of chinmo for the larval growth of SGs and wing discs. (A) DAPI-stained SGs from control (*fkhGal4*) and *UASBr-C*^*RNAi*^; *UASchinmo*^*RNAi*^ L3W larvae. In the absence of *chinmo* and Br-C, SGs did not grow. (B-D) Comparison of the relative size of SGs (n = 10 for each genotype) (B), DAPI intensity (n = 50 for each genotype) (C), and nucleic size of SGs (n = 30 for each genotype) (D) of control and *UASBr-C*^*RNAi*^; *UASchinmo*^*RNAi*^ L3W larvae. (E) Relative expression of SG secretion genes in control and *UASBr-C*^*RNAi*^; *UASchinmo*^*RNAi*^ L3W larvae measured by qRT-PCR. Transcript abundance values were normalised against the *Rpl32* transcript. Fold changes were relative to the expression of the control, arbitrarily set to 1 (dashed black line). Error bars in B and C indicate the SEM (n = 5-8). Asterisks in B-D indicate differences statistically significant at ∗ ∗ ∗*p ≤ 0*.*001*. (F) Expression of Chinmo, Br-C and Wg (E) in wing discs of *UASBr-C*^*RNAi*^*;UASchinmo*^*RNAi*^ L3W larvae. Wing discs labelled to visualise the esg domain (GFP in green). In the absence of chinmo and Br-C, wing discs grow normally and express Wg correctly. Scale bars represent 50 µm. **Figure 5—Source Data 3** **Numerical data for Figure 5B-E**.

### Down-regulation of *chinmo* is required during metamorphosis

The functional and expression data reported above show that Chinmo acts as a larval specifier in *Drosophila*. From this, we could infer that its absence by the end of larval development is required first for the transition to the prepupa, and then to allow terminal adult differentiation during the pupal period. If this were the case, then, the maintenance of high levels of *chinmo* during late L3 would interfere with the larva-pupal transition. To test this possibility, we maintained high levels of *chinmo* in late L3 wing discs using the Gal4/Gal80^ts^ system. Consistent with our hypothesis, overexpressing *chinmo* from early L3 in the anterior compartment of the disc using the *cubitus interruptus ciGal4* driver abolished *Br-C* expression and induced apoptosis in this compartment at late L3 as revealed by the high expression of the effector caspase Dcp-1 (Figure 6A). As a result, the size of the anterior compartment was dramatically reduced, and the expression of patterning genes such as *ct* was halted (Figure 6B). The impairment of *ct* expression was not due to the death of the tissue, but to a specific response to the sustained expression of *chinmo* or to depletion of *Br-C*, as its expression was still not detected in wing discs overexpressing both *chinmo* and the p35 inhibitor of effector caspases (Hay et al., 1994) (Figure 6C and D).

**Figure 6.**
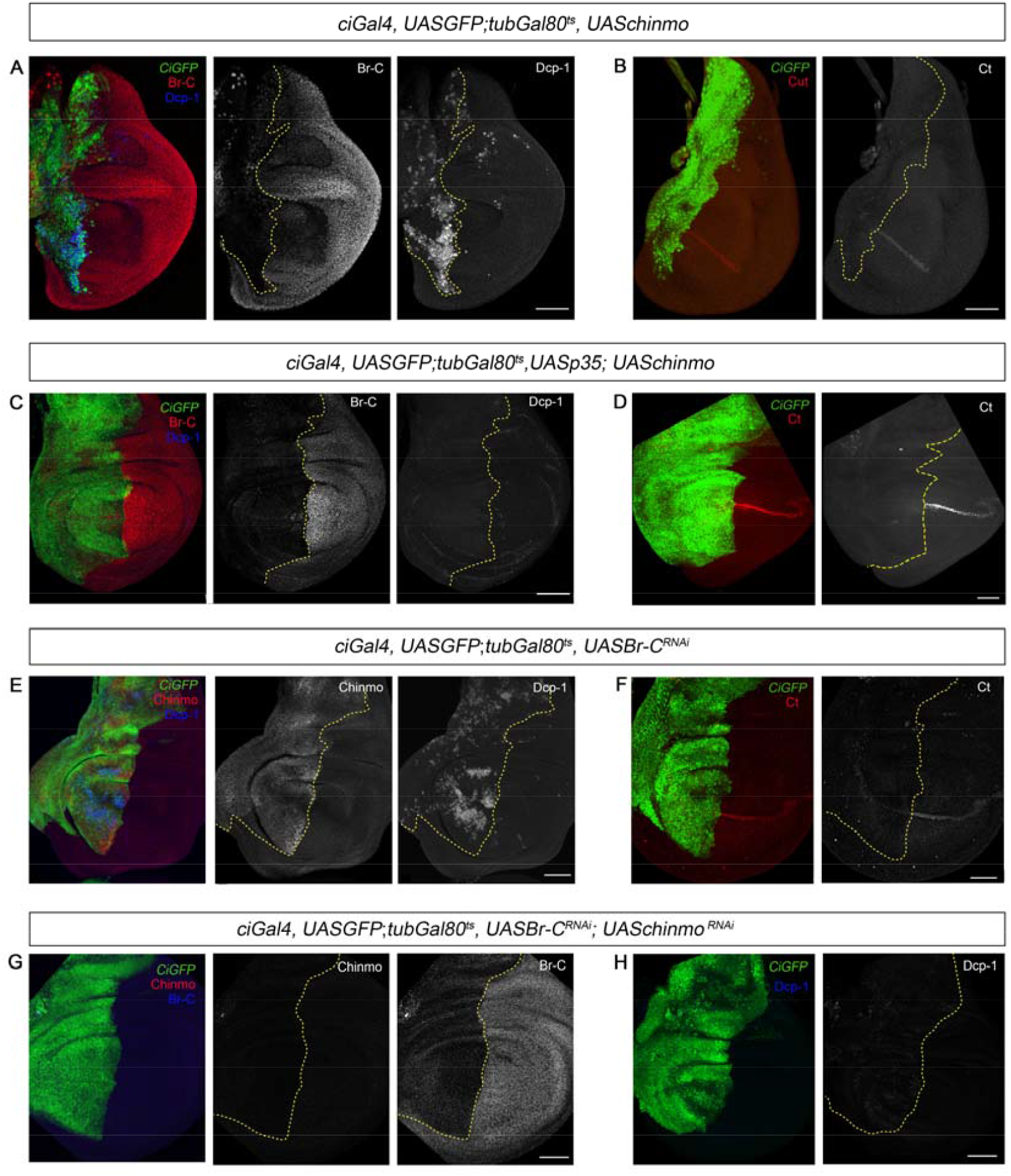
Chinmo depletion during late L3 is required for proper larva to pupa transition. (A-H) Images of wing imaginal discs from L3W larvae. The indicated constructs were expressed under the control of the *ciGal4* driver. Overexpression or depletion of the transgenes was activated in early L3 larvae and analyzed at the L3W stage. An UAS-GFP construct was used to mark the anterior region of the disc where the transgenes were induced or repressed (green). (A) Overexpression of *chinmo* repressed Br-C, induced Dcp-1 and (B) depleted Cut. (C) Overexpression of *chinmo* together with *p35* repressed Br-C and blocked Dcp-1, but fails to restore normal expression of Ct (D). (E) Depletion of *Br-C* induced Chinmo and Dcp-1 and (F) repressed Ct. (G) In double depletion of *Br-C* and *chinmo* (H), Dcp-1 was not detected. Scale bars represent 50 µm.

An alternative way to maintain high levels of *chinmo* in late L3 is by depleting *Br-C*, a well-known repressor of *chinmo* from mid-L3 (Narbonne-Reveau and Maurange, 2019). Therefore, we knocked down *Br-C* in the anterior compartment of the wing disc. As expected, Chinmo levels remained high in this compartment of late L3 wing disc, with a concomitant strong Dcp-1 staining and impairment of *ct* expression (Figure 6E and F). Importantly, simultaneous depletion of *chinmo* and *Br-C* from early L3 did not lead to an increase in apoptosis (Figure 6G) nor alter the expression of patterning genes (Figure 5F), which indicates that tissue death at the end of the larval period is due to sustained expression of Chinmo rather than the absence of Br-C. Altogether, these results confirm that the transition from larva to pupa must take place in the absence of the larval specifier Chinmo.

Next, we analyzed whether lack of *chinmo* is also important during the pupal period for the E93-dependent development of the adult. To this end, we used the thermo-sensitive system to overexpress *chinmo* in the anterior part of the wing specifically during the pupal stage. This ectopic expression of *chinmo* led to a marked decrease in E93 protein levels (Figure 7A). As a result, the anterior compartment of the wing was strongly undifferentiated, a phenotype reminiscent of that observed in E93-depleted wings (Ureña et al., 2014, 2016) (Figure 7B). Taken together, our results show that *chinmo* must be downregulated during the initiation and throughout the metamorphic transition to allow the sequential expression of the pupal specifier *Br-C* and the adult specifier *E93*.

**Figure 7.**
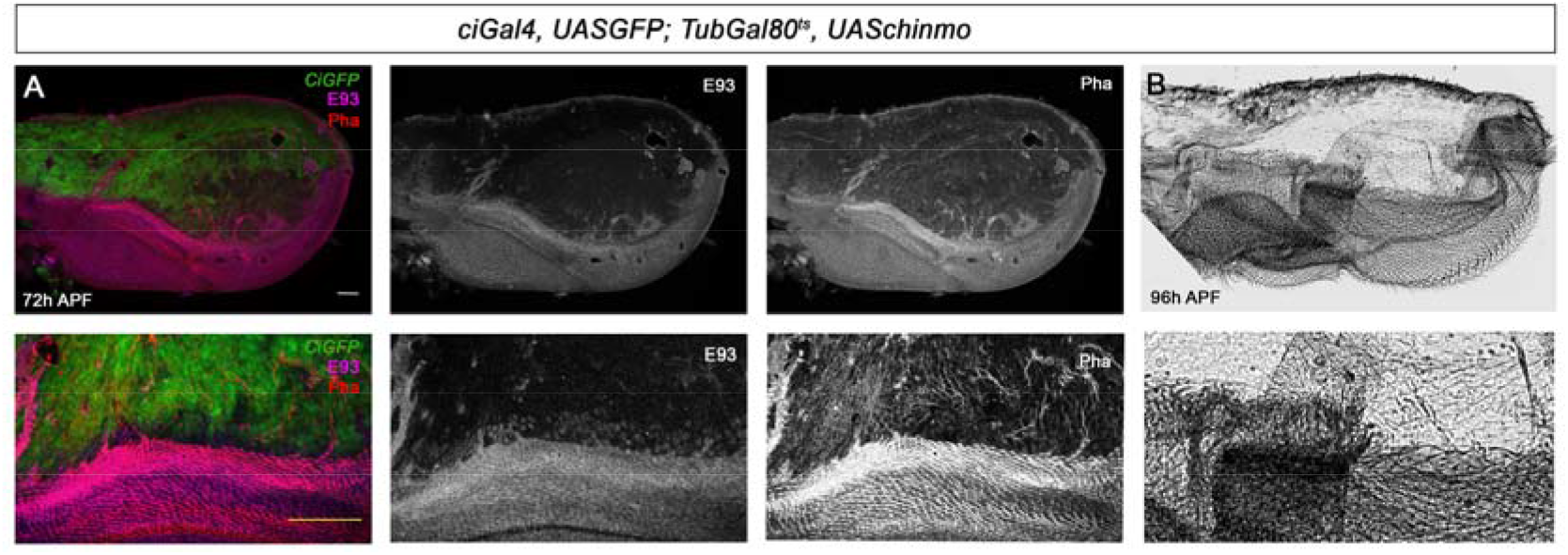
Presence of Chinmo during pupal development blocks adult differentiation. (A) Overexpression of *chinmo* in the anterior part of the pupal wing at 72 h after pupa formation (APF) using *ciGal4* driver represses *E93* expression and produced alterations in Phalloidin (Pha) pattern. (B) Cuticle preparation of a pupal wing at 96 h APF expressing *chinmo* under the control of the *CiGal4* driver. Bottom panels are magnifications from upper images. The scale bars represent 50 μm (top panels) and 100 μm (bottom panels).

### Antagonistic effects of *chinmo*, and *Br-C*/*E93* in tumour growth

Chinmo and Br-C belong to the extended family of BTB-ZF transcription factors, which are not restricted to insects. In humans, many such factors have been implicated in cancer, where they have opposing effects, from oncogenic to tumour suppressor functions (Siggs and Beutler, 2012). However, while overexpression of *Drosophila chinmo* has been found to cooperate with Ras or Notch to trigger massive tumour overgrowth (Doggett et al., 2015), changes in *Drosophila Br-C* expression have not been associated with any effect on tumorigenesis. Since our results described inhere, and those from other labs (Narbonne-Reveau and Maurange, 2019) indicate that *chinmo* and *Br-C* have antagonistic effects in terms of proliferation vs. differentiation, we addressed whether these opposite features might also be associated with pro-oncogenic or tumour suppressor properties, respectively. To test this notion, we resorted to the well-defined tumorigenesis model in *Drosophila* generated by the depletion of cell polarity genes such as *lgl* (Froldi et al., 2008; Gong et al., 2021). To suppress *lgl* and create an oncogenic sensitised background (Figure 8A, B and G), we used two *UASlgl*^*RNAi*^ constructs recombined on the third chromosome to drive their expression by *nubbinGal4* (*nubGal4*) in the imaginal wing disc pouch.

**Figure 8.**
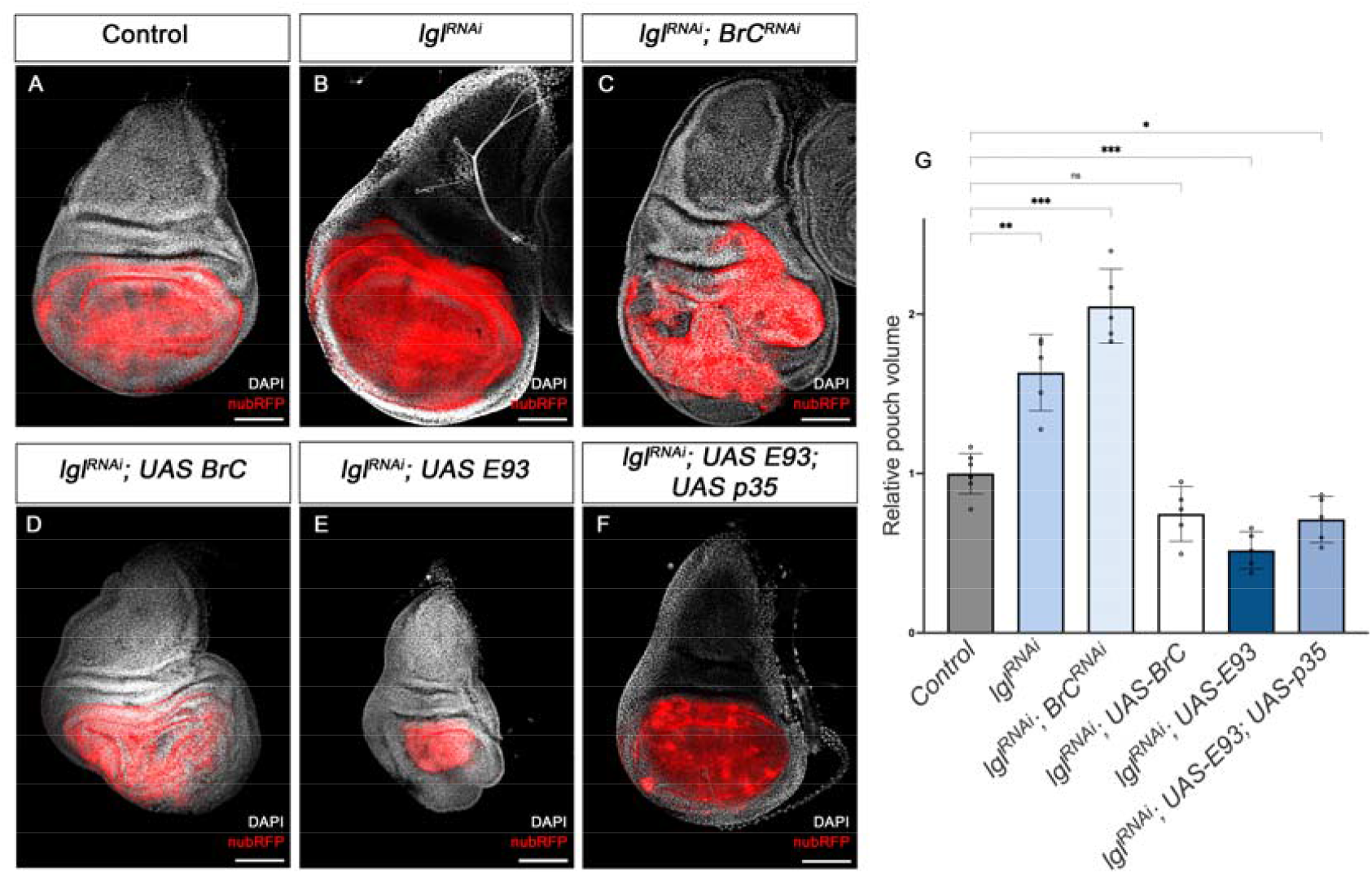
Pro-oncogene and tumour suppression action of chinmo and Br-C, respectively. (A-F) Confocal images of L3 wing imaginal discs. The indicated constructs were expressed under the control of the *nubGal4* driver. A *UASRFP* construct was used to mark the pouch region of the disc where the transgenes were induced (magenta). Nuclei were labelled with DAPI (grey). Scale bars at 100 μm. (G) Volumetric quantification of the RFP-positive area of the wing discs for the indicated groups. The pouch volumes were normalised to the mean of the control. Welch’s ANOVA (*p<0*.*0001*) followed by Dunnett’s T3 post hoc tests (*\* p<0*.*05, ** p<0*.*01, *** p<0*.*001*). **Figure 8—Source Data 4** **Numerical data for Figure 8G**.

Interestingly, RNAi-mediated depletion of *Br-C* in the wing discs in the downregulated *lgl* background resulted in an increase in the mean wing pouch volume compared to the downregulation of *lgl* alone (Figure 8B, C and G). Consistently, overexpression of *Br-C* in the same *lgl* background had the opposite effect, reducing the size of the *lgl*-induced overgrowth, thereby confirming that *Br-C* expression elicits tumour suppressor activity (Figure 8D and G). Given that *E93* has a similar pro-differentiation role to that of *Br-C*, we also examined whether *E93* also exerts tumour suppressor activity. We found that overexpression of *E93* also reduced the size of *lgl* overgrowth (Figure 8E, G). However, as *E93* overexpression triggers cell death in some tissues (Pahl et al., 2019), we wanted to assess whether the reduction of the pouch region in this case was caused by apoptosis induction. When *E93* overexpression was combined with the p35 inhibitor of apoptosis, we still observed a reduction in the size of the wing pouch in the *lgl-*sensitised background (Figure 8F, G). Thus, even in this regard, *chinmo* plays an opposite function than Br-C and E93; while *chinmo* has pro-oncogenic features, *Br-C* and *E93* act as tumour suppressors.

### Role of chinmo in hemimetabolous development

As full metamorphosis is an evolutionary acquisition of holometabolous insects from hemimetabolous ancestors with no pupal stage (Truman, 2019), we sought to determine whether the role of chinmo as a larval specifier was also present in hemimetabolous insects. To this end, we used the German cockroach *Blattella germanica* as a model for hemimetabolous development. *Blatella* goes through six juvenile nymphal instars (N1-N6) before developing into an adult. Metamorphosis takes place during N6 and is restricted to the transformation of the wing primordia into functional wings, the attainment of functional genitalia, and changes in cuticle pigmentation (Ureña et al., 2014). A detailed Tblastn search in the *Blatella* genome database revealed the presence of a *chinmo* orthologue (*Bg-chinmo*).

To study *Bg-chinmo*, we first examined its expression during the life cycle of *Blattella*. We found that it is highly expressed in embryos and decreases dramatically thereafter during nymphal development (Figure 9A). In order to analyze the function of the relative low levels of *Bg-chinmo* during postembryonic satges, we analysed the function of *Bg-Chinmo* by systemic injection of dsRNAs into newly emerged N4 instar. Specimens injected with dsMock were used as negative controls (Control animals). Importantly, whereas Control larvae underwent two larval molts before initiating metamorphosis at the end of the N6 stage, 43% of *Bg-chinmo*^*RNAi*^ animals underwent only one nymphal molt before molting to an early adult after the N5 stage (Figure 9B-E). Precocious *chinmo*-depleted adults were smaller than control counterparts as they skipped a nymphal stage. However, they presented all the external characteristics of an adult, namely functional hind- and fore-wings, adult cerci, and adult-specific cuticle pigmentation. Altogether, these results suggest that the role of Chinmo as juvenile specifier seems to be conserved in hemimetabolous insects, thereby indicating that its developmental function precedes the hemimetabolous-holometabolous split.

**Figure 9.**
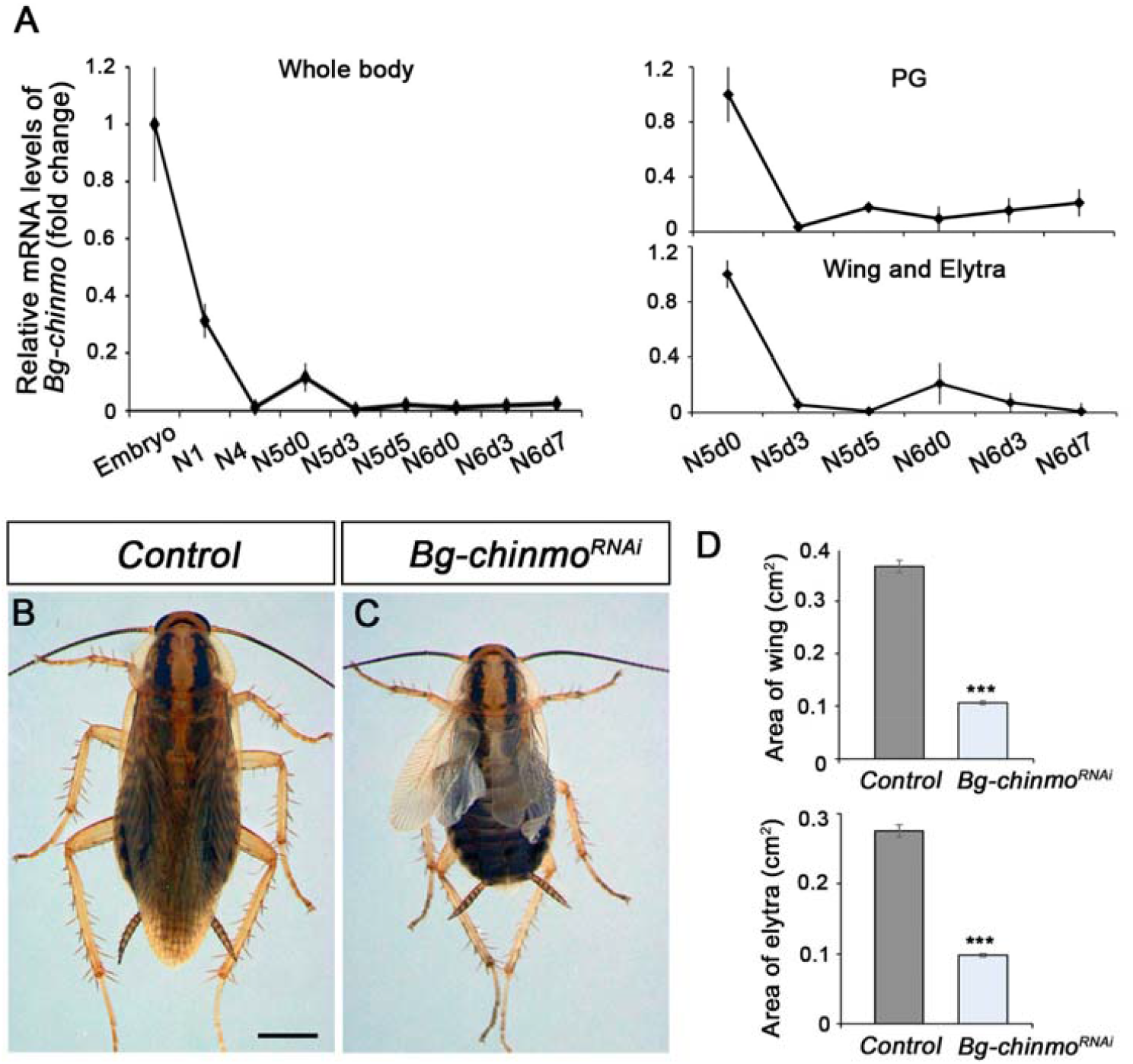
Depletion of *chinmo* in *B. germanica* promotes premature adulthood. (A) *Bg-chinmo* mRNA levels measured by qRT-PCR from embryo to the last nymphal stage (N6) in whole body, prothoracic gland (PG), and wings and elytra. Transcript abundance values were normalised against the *Rpl32* transcript. Fold changes were relative to the expression of embryo (for whole body) or N5d0 (for PG and wings and elytra), arbitrarily set to 1. Error bars indicate the SEM (n = 3-5). (B-C) Newly moulted N4 nymphs of *B. germanica* were injected with *dsMock* (*Control*) or *dschinmo* (*Bg-chinmo*^*RNAi*^) and left until adulthood. (B) Dorsal view of adult Control, and (C) Premature adult *Bg-chinmo*^*RNAi*^. (D) Quantification of wing and elytra areas (cm^2^) of adult Control and *Bg-chinmo*^*RNAi*^ premature adults. Statistical significance was calculated using t-test (∗ ∗ ∗*p ≤ 0*.*001*). The scale bar represents 2 mm. **Figure 9—Source Data 5** **Numerical data for Figure 9A and D**.

In summary, we identified Chinmo as a new member of the MGN acting as a general larval specifier, as recently proposed by Truman and Riddiford (Truman and Riddiford, 2022). Together with a number of previous reports (reviewed in (Martín et al., 2021)), our results show that the temporal expression of Chinmo, Br-C and E93 determine the tissue adquisition of gradual differentiation features from the juvenile to the adult to generate the distinct organs. Whereas Chinmo maintains cells in an undifferentiated state, Br-C and E93 induce progressively the differentiation program. This effect has already been shown in the central nervous system where early-born neurons are characterized by the expression of Chinmo, whereas smaller late-born neurons are marked by expression of Br-C (Maurange et al., 2008). Similarly, the *chinmo*-to-*Br-C* transition in *Drosophila* has been associated with the loss of the regenerative potential of imaginal cells (Narbonne-Reveau and Maurange, 2019). The fact that Br-C and E93 acts a tumor supressor in a overproliferative backgroung supports this idea.

Finally we found that the role of chinmo as larval specifier is conserved in hemimetabolous insects. Since hemimetabolous insects do not undergo the intermediate pupal stage, the transition from juvenile to adult, therefore, relies exclusively on the shift from Chinmo to E93 during the last nymphal stage, with Kr-h1 also involved in preventing metamorphosis through the repression of *E93* (Ureña et al., 2016). Interestingly, Br-C in most hemimetabolans has been shown to exert no role during development (Erezyilmaz et al., 2006; Konopova et al., 2011), suggesting that this factor has been co-opted as a pupal specifier during the arise of holometabolous insects.

## Materials and Methods Fly strains

All fly stocks were reared at 25°C on standard flour/agar Drosophila media. The Gal4/UAS system was used to drive the expression of transgenes at 29°C. Gal4/Gal80ts system was used for conditional activation. In these experiments, crosses were kept at 18 until L2 or L3-late molt and then shifted to 29ºC for conditional induction. The following strains used in this study were provided by the Bloomington Drosophila Stock Center (BDSC): *fkhGal4* (#78060); *ActGal4* (#3954); *TubGal80ts* (#7016), *UASchinmo*^*RNAi*^ (#26777); *UASchinmo* (#50740); *UASBr-C*^*RNAi*^ (#51378); *UASP35* (#5072); *UAS-myr-mRFP* (#7118) and *UAS-mCD8::GFP* (#32186) were used to follow the GAL4 driver activity. Two lines of *UASlgl*^*RNAi*^ (#51247 and #51249) were obtained from Vienna Drosophila RNAi Center (VDRC). *nubGal4* (Calleja et al., 1996), *esgGal4UASGFP* (Jiang et al., 2009) and *ciGal4* (Croker et al., 2006) were used to drive the expression of different constructs in the wing disc. Crosses to *CantonS* line were used as control.

### Blattella germanica

Specimens of B. germanica were obtained from a colony reared in the dark at 30 ± 1 °C and 60–70% relative humidity. Cockroaches undergo hemimetabolous development, where growth and maturation take place gradually and simultaneously during a series of nymphal instars. In our rearing conditions, *B. germanica* undergoes six nymphal instars (N1–N6) before molting into the adult. All dissections and tissue sampling were carried out on carbon dioxide anesthetized specimens.

### Immunohistochemistry

For fluorescent imaging, SGs, wing discs from different juvenile stages and pupal wings were dissected in 1XPhosphate-Buffered Saline (PBS) and fixed in 4% formaldehyde for 20 min at RT. The tissues were rinsed in 0.1% Triton X-100 (PBST) or 0.3% PBST in pupal wings for 1h and incubated at 4°C with primary antibodies diluted in PBST overnight. After incubation with primary antibodies, the tissues were washed with PBST (3 × 10min washes) and incubated with adequate combinations of secondary antibodies (Alexa Conjugated dyes 488, 555, 647, Life Technologies, 1:500) for 2h at RT, followed by 3 × 10min washes with PBST, and then rinsed with PBS before mounting in Vectashield with DAPI (Vector Laboratories, H1200) for image acquisition. The following primary antibodies were used at indicated dilution: rat anti-Chinmo (1:500, N, Sokol), mouse anti-Cut (1:200, Developmental Studies Hybridoma Bank (DSHB) #2B10), mouse anti-Wg (1:200, DSHB #4D4), mouse anti Br-C core (1:250 DSHB #25E9.D7), rabbit anti-cleaved Dcp-1 (1:100, Cell Signaling #9578) and rabbit anti-E93 (1:50, this work).

### Antibody generation

A peptide corresponding to the 23 residues (GRRAYSEEDLSRALQDVVANKL) of E93 was coupled to KLH and was injected into rabbits. Polyclonal antisera were affinity-purified and were found to be specific for E93, by Western blotting and by immuno-fluorescence.

### RNA extraction and quantitative real-time reverse transcriptase polymerase chain reaction (qRT-PCR)

Total RNA was isolated with the GenElute Mammalian Total RNA kit (Sigma), DNAse treated (Promega) and reverse transcribed with Superscript II reverse transcriptase (Invitrogen) and random hexamers (Promega). In the case of *Drosophila*, cDNA was obtained from whole larvae (CantonS) or L3 wandering SGs. *Blattella germanica* cDNAs were obtained from whole nymphs or wings and PG of different juvenile instars. Relative transcripts levels were determined by real-time PCR (qPCR), using iTaq Universal SYBR Green Supermix (Bio-Rad). To standardize the qPCR inputs, a master mix that contained iTaq Universal SYBR Green PCR Supermix and forward and reverse primers was prepared (final concentration: 100nM/qPCR). The qPCR experiments were conducted with the same quantity of tissue equivalent input for all treatments and each sample was run in duplicate using 2μl of cDNA per reaction. All the samples were analyzed on the iCycler iQ Real Time PCR Detection System (Bio-Rad). RNA expression was calculated in relation to the expression of *DmRpl32* or *BgActin5C*. Primers sequences for qPCR analyses were (Duan et al., 2020):

*DmChinmo*-F: 5’ AGTTCTGCCTCAAATGGAACAG ‘3

*DmChinmo*-R: 5’ CGCAGGATAATATGACATCGGC ‘3

*Dm*Sgs1-F: 5’CCCAATCCCGTGTGGCCCTG ‘3

*Dm*Sgs1-R: 5’ GTGATGGCAACGGCGGTGGT ‘3

*Dm*Sgs3-F: 5’ TGCTACCGCCCTAGCGAGCA ‘3

*Dm*Sgs3-R: 5’ GTGCACGGAGGTTGCGTGGT ‘3

*Dm*Sgs4-F: 5’ ACGCATCAAGCGACACCGCA ‘3

*Dm*Sgs4-R: 5’TCCTCCACCGCCCGATTCGT ‘3

*Dm*Sgs7-F: 5’ CGCAGTCACCATCATCGCTTGC ‘3

*Dm*Sgs7-R: 5’ACAGCCCGTGCAGGCCTTTC ‘3

*Dm*Sgs8-F: 5’ AGCTGCTCGTTGTCGCCGTC ‘3

*Dm*Sgs8-R: 5’ GCGGAACACCCAGGACACGG ‘3

*DmRpL32*-F: 5’CAAGAAGTTCCTGGTGCACAA’3

*DmRpL32*-R: 5’AAACGCGGTTCTGCATGAG’3

*BgChinmo-F*: 5’ CAGCACCACTATGTCCAAGTG ‘3

*BgChinmo-R*: 5’ CAGGAAACTGGAGAGGCTTTC ‘3

*BgActin5C-F*: 5′-AGCTTCCTGATGGTCAGGTGA-3′

*BgActin5C-R*: 5′-TGTCGGCAATTCCAGGGTACATGGT-3′

## RNA interference (RNAi)

RNAi in vivo in nymphs was performed as previously described (Cruz et al., 2007; Martín et al., 2006). A dose of 1 µl (4–8 µg/µl) of the dsRNA solution was injected into the abdomen of newly antepenultimate (N4d0) instar nymphs, and left until analysed. To promote the RNAi effect, the same dose of dsRNAs was reapplied to all treated animals after three days (N4d3) from the first injection. Control dsRNA consisted of a non-coding sequence from the pSTBlue-1 vector (dsControl). The primers used to generate templates via PCR for transcription of the dsRNA were:

*BgChinmo-F*: 5’CAGCACCACTATGTCCAAGTG’3

*BgChinmo-R*: 5’GAGTCCTGCATGGCTTCGGA’3

## Imaging acquisition and analysis

Images were obtained with the Leica TCS SP5 and the Zeiss LSM880 confocal microscopes. The same imaging acquisition parameters were used for all the comparative analyses. Images were processed with the Imaris Software (Oxford Instruments), Fiji or Photoshop CS4 (Adobe). For DNA quantification and nuclear size of SGs, DNA staining intensity in the SG cells was obtained from z stacked images every 0.25 μm of DAPI stained L3 larvae. Image analysis was performed using Fiji. The volume of the wing pouch region was measured in Imaris software (Oxford Instruments). Adult flies, nymphal parts and adult cockroach images were acquired using AxioImager.Z1 (ApoTome 213 System, Zeiss) microscope, and images were subsequently processed using Photoshop CS4 (Adobe).

## Statistical analysis

Statistical analysis and graphical representations were performed in GraphPad Prism 9 software. All experiments were performed with at least three biological replicates. Two-tailed Student’s test and Welch’s ANOVA followed by Dunnett’s T3 post hoc tests were used to determine significant differences.

## Acknowledgements

The ICTS “NANOBIOSIS”, and particularly the Custom Antibody Service (CAbS, IQAC-CSIC, CIBER-BBN), is acknowledged for the assistance and support related to the E93 antibody used in this work. This project is supported by the Spanish MINECO (grant PGC2018-098427-B-I00 and PID2021-125661NB-100 to D.M., and X.F-M. and grant PGC2018-094254-B-100 to J.C) and by the Catalan Government (2017-SGR 1030 to D.M. and X.F-M and trhought BIST to J.C.). The research has also benefited from FEDER funds. S.C. is a recipient of a Juan de la Cierva contract from the MINECO.

## Supplementary Information

**Supplementary Figure 1.**
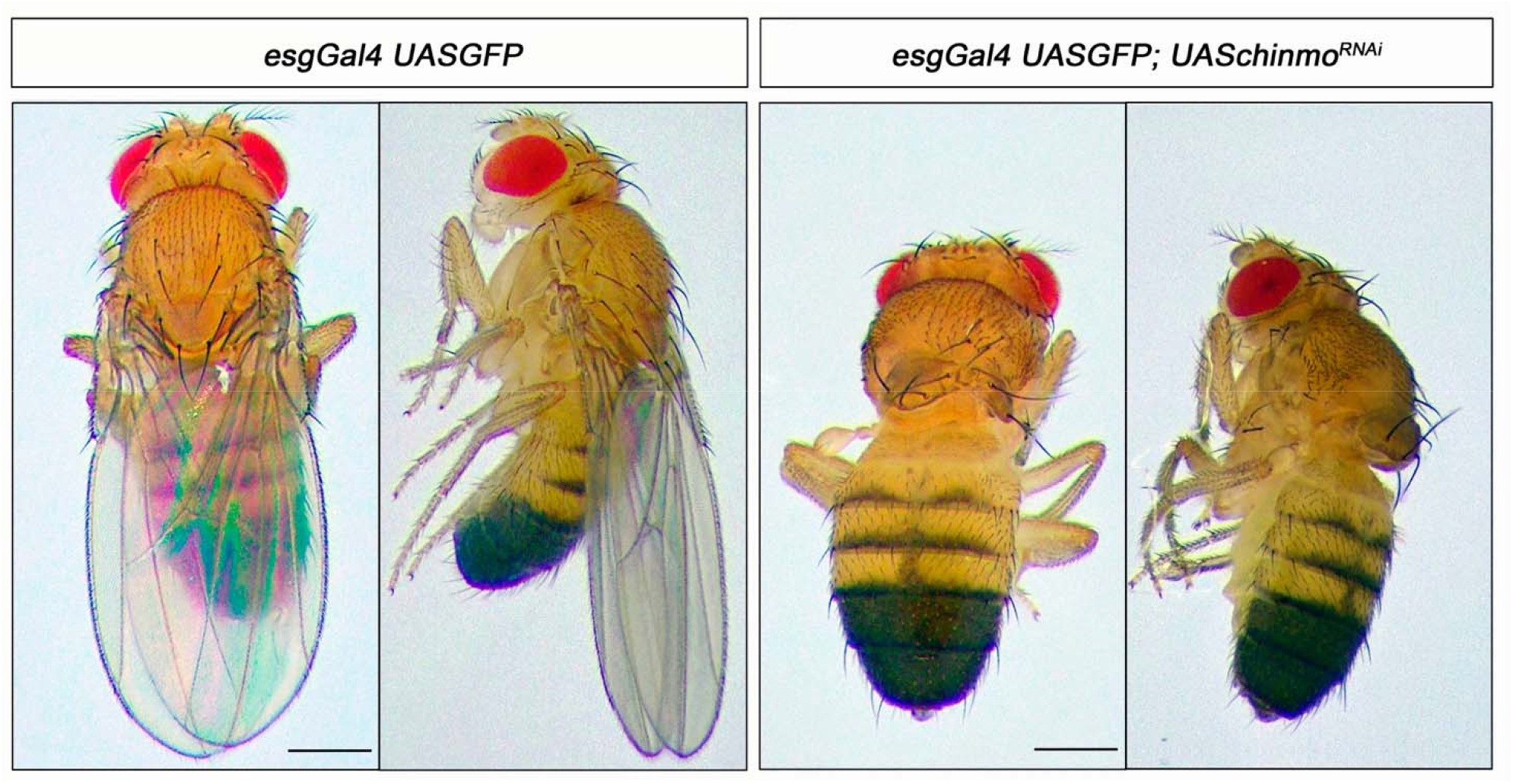
Chinmo is required for wing development during the larval period. Dorsal and lateral view of control (right panel) and *UASchinmo*^*RNAi*^ (left panel) adult flies. In the absence of Chinmo, flies emerged without wings. Scale bar represents 1 mm.

**Supplementary Figure 2.**
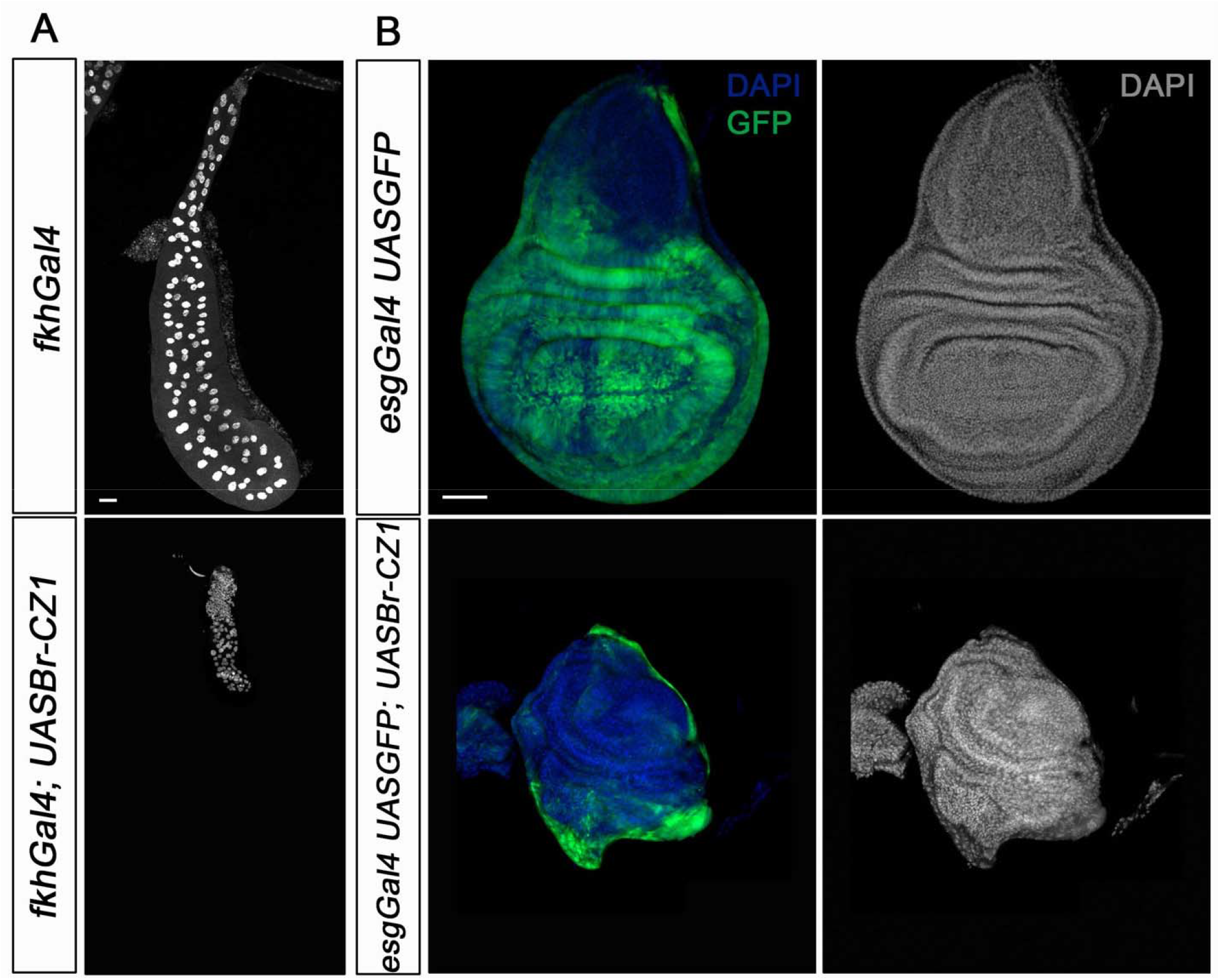
Overexpression of Br-CZ1 phenocopies *chinmo* loss of function in SGs and wing discs. (A) DAPI-stained SGs from control (*fkhGal4*) and *UASBr-CZ1* in L3W larvae. Overexpression of Br-CZ1 impairs SGs grow. (B) Wing discs of control (*esgGal4*) and *UASBr-CZ1* L3W larvae. Wing discs were labelled to visualize the *esg* domain (GFP in green) and nuclei (DAPI). Overexpression of Br-CZ1 in the *esg* domain abolishes wing development. Scale bar represents 50 µm in all panels.

